# Water salinity does not affect acute thermal tolerance (CT_max_) in zebrafish (*Danio rerio*)

**DOI:** 10.1101/2022.08.02.502531

**Authors:** Eirik R. Åsheim, Anna H. Andreassen, Rachael Morgan, Mireia Silvestre, Fredrik Jutfelt

## Abstract

Tolerance against acute warming is an essential trait that can determine how organisms cope during heatwaves, yet the mechanisms underlying it remain elusive. Water salinity has previously been shown to modulate thermal tolerance and may therefore provide clues towards these limiting mechanisms. Here we tested whether short (2 hours) and long (10 days) term exposure to different water salinities (0-5 ppt) affected acute thermal tolerance in zebrafish (N=269). We found that water salinity did not affect the thermal tolerance of zebrafish at either time point, indicating that salinity does not affect the mechanism limiting acute thermal tolerance limits in zebrafish. We did, however, observe unexpected behaviour during the CT_max_ test in a subset of fish in the highest salinity treatment after 10 days (3 ppt), indicating some form of salinity-driven disturbance during warming.

## INTRODUCTION

With climate warming, average water temperatures are increasing, and in many places, temperature extremes are increasing in frequency, duration and maximum temperatures (Seneviratne et al., 2014). In some aquatic habitats, such heat waves may push populations to, or above, their acute upper thermal tolerance limits. Such situations may already be occurring in some coral reefs (Genin et al., 2020), and freshwater systems (Morgan et al., 2019; Wegner et al., 2008). As the increasingly severe heatwaves will increase the impacts from extreme temperatures, there is increased research interest in understanding the mechanisms behind warming tolerance.

The most commonly used measure of acute upper thermal limits in aquatic ectotherms is the critical thermal maximum test (CT_max_), where the water temperature is gradually increased until the animal displays a loss of balance or locomotor function (Becker & Genoway, 1979; Lutterschmidt & Hutchison, 1997; Morgan et al., 2018). The temperature at loss of equilibrium (LOE) is commonly used as the endpoint for actively swimming fishes. However, the mechanisms causing the LOE are unknown. With a lack of understanding of the underlying physiology, it is also difficult to predict how temperature may interact with other factors.

One such other factor is water salinity. Salinity stress can occur in aquatic environments through processes such as migration between water bodies, evaporation, ice melt, changes in stratification, and precipitation. As climate change affects weather patterns, some fish populations will likely experience some form of osmotic stress, either through increased or decreased salinity.

Examining how water salinity affects thermal tolerance can be useful for predicting interactions of factors during heat waves in nature. If water salinity modulates thermal tolerance limits, then differences in precipitation or evaporation may affect the thermal tolerance of some populations in nature. Additionally, modulation of thermal tolerance by other factors may give information on which physiological mechanisms are controlling these traits. Such interactions could for example indicate if plasma osmolality and ion composition are involved in the mechanisms controlling thermal tolerance.

There have been a few indications in the recent literature suggesting that water salinity can modulate thermal tolerance, but there has been no consistent direction of effects with salinity manipulation (Figure 1). Some studies report a reduction in thermal tolerance with increased salinity (Morgan et al., 2019; Shaughnessy & McCormick, 2018), some show increased thermal tolerance with increasing salinity (King & Sardella, 2017; Metzger et al., 2016), some show a peak thermal tolerance around some optimal salinity (Haney & Walsh, 2003; Sardella et al., 2008), whilst others report no effect (Davis et al., 2019; Hines et al., 2019) (Suppl. Table 1). Several different mechanisms have been suggested to be responsible where CT_max_ has been affected, such as oxygen availability (via changes in blood hematocrit and lactate)(Shaughnessy & McCormick, 2018), expression of heat shock proteins (Metzger et al., 2016), and changes in gill permeability (Haney & Walsh, 2003) However, none of these suggested mechanisms have so far been confirmed.

**Figure 1.**
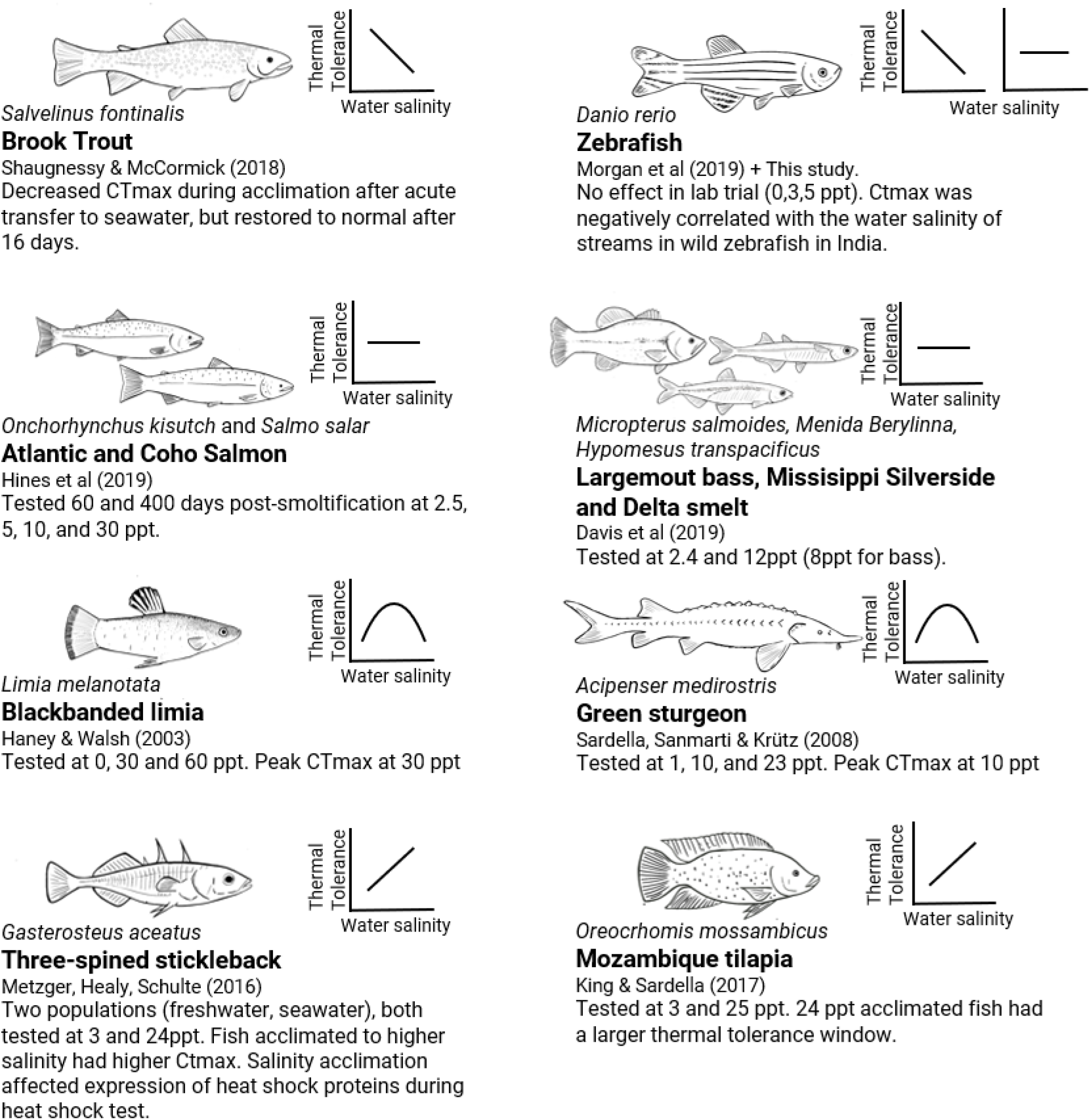
Summary of currently available reports from studies testing CT_max_ under various water salinity treatments in different fish species. The small graphs indicate the general, simplified pattern of the change in CT_max_ as salinity increases (Increased, decreased, no change, or peaked). Different species of fish and different experimental settings give different effects of salinity on thermal tolerance.

A field study by Morgan et al. (2019) suggested a reduction in CT_max_ with higher salinities. In that study, CT_max_ of wild zebrafish (*Danio rerio*) correlated negatively with water salinity, with fish from higher salinity streams (measured as conductivity) showing lower thermal tolerance (Fig. 1a). It was, however, not clear if that effect was due to water salinity or some other confounding factor of the sampled river systems. This curious observation that stream salinity appeared to affect thermal tolerance, together with the extensive amount of research tools available for the species, suggested that zebrafish could make a good study system for research on the effects of salinity on thermal tolerance.

To test the findings by Morgan et al. (2019), to explore the zebrafish as a model species for interactions between salinity and warming tolerance, and to build on the existing knowledge-body on this topic, we designed an experiment testing CT_max_ of zebrafish after exposure to different salinity treatments. We were also interested in knowing whether a potential effect is immediate (e.g. due to acutely disturbed plasma ion concentrations or altered electrochemical ion gradients), and if such an effect is modulated by longer exposures and acclimation to the water salinity treatments. Therefore, we tested fish at two different time points; after two hours and after ten days of salinity exposure.

## METHODS

### Fish rearing and experimental design

Fish used in this study were randomly selected from the first filial generation of wild-caught zebrafish from West Bengal, India (Morgan et al., 2019, 2022). For the duration of the experiment, fish were held in 45 L glass aquaria (50 × 30 × 30cm) filled to 35 L, at 26°C, and under a 12/12 day/night cycle. In each tank, an ornamental aquarium plastic plant was used to enrich the tank environment, and water was filtrated and aerated using two bacteria-seeded air-lift sponge filters. Fish were fed ad-libitum twice a day using TetraPro energy flakes (Tetra ®, Blacksburg, VA, USA).

We divided the fish into eight experimental tanks, having two duplicate tanks per treatment, with at least 30 individuals in each tank. The total number of fish was 269, with 67 fish at 0 ppt, 69 at 0.5 ppt, 67 at 1 ppt, and 66 at 5 ppt. Different levels of water salinity were achieved by adding natural sea salt (GC Rieber salt A/S, 98.0% NaCl) to aerated and carbon-filtered tap water and measured to 0.1 ppt accuracy with a digital water quality meter (YSI ProDSS, YSI inc, USA) using a conductivity measurement probe (ProDSS ODO/CT, YSI Inc, USA). The fish were kept for three days to habituate to their experimental tanks at 0.5 ppt before the salinity treatments started.

### Salinity exposures

The experimental design included four treatment groups of different salinity, and each treatment had two duplicate tanks. Prior to the experiment, the fish were kept at 0.5 ppt, which is considered optimal holding conditions for laboratory zebrafish (Lawrence, 2007). One group was kept at this concentration as a control group. A group kept at 1 ppt served as a “high” treatment, and a group kept at of 5 ppt served as a “very-high” treatment. To see if hypoosmotic stress could also affect thermal tolerance, we included a 0 ppt group using millipore-filtered water. All fish were acutely exposed to the salinity treatments. Salinity was changed by moving the fish to a temporary tank while replacing the water in the experimental tank with water at the target salinity.

Zebrafish is a freshwater species, and the observations available on the salinities they experience in the wild indicate that they live at low salinities around 0.1-0.6 ppt (Morgan et al., 2019; Spence et al., 2006). However, a laboratory experiment testing acute salinity tolerance in zebrafish found no effect on metabolism or nitrogen excretion at salinities as high as 20 ppt. Knowing this, we chose 5 ppt as the highest salinity treatment for our experiment as we wanted a strong positive control in case we found no effect, while staying far below what we knew these fish could survive acutely. The fish were regularly monitored after the initial change of salinity, looking for any change in behaviour or activity that could indicate distress. After two days day at the 5 ppt treatment, some fish displayed signs of inactivity (e.g. lying still on the bottom), which indicated that the 5 ppt salinity treatment was too high. Therefore, the salinity concentration was decreased to 3 ppt for the remaining ten-day treatment. This is still 6 times higher than the recommended salinity for zebrafish (Lawrence, 2007), and 50 times higher than the 0.06±0.02 ppt (±SD) salinity measured in their native Indian streams (Morgan et al., 2019; Sundin et al., 2019).

The fish were tested for acute thermal tolerance at two different time points. Half of each treatment group (each group n=28-39) was tested for CT_max_ 130-150 minutes after the start of salinity exposure, and the other half after ten days of salinity exposure.

### Testing of acute thermal tolerance (CT_max_)

The CT_max_ tests were conducted as in Morgan et al. (2018). Briefly, the CT_max_ method consists of a plexiglass tank (33 ⋅23 ⋅24 cm) with two compartments separated by a mesh: one heating compartment and one fish arena. The heating compartment contained a 300 W heater coil placed within a custom-made heating chamber and attached to a water pump to maintain a homogenous temperature within the tank (Morgan et al., 2018). The tank was filled with 9.9 L of 26°C water at the corresponding salinity concentration for each treatment. A group of 7–11 fish were placed in the arena simultaneously and the temperature was gradually increased by a rate of 0.3°C per minute.

The temperature at which the fish lost equilibrium (defined as disorganised swimming for three seconds, usually rolling during swimming) was recorded with 0.1°C precision for each individual, using a high precision thermometer (Testo-112, Testo, Titisee-Neustadt, Germany). Other behavioural abnormalities were noted but not quantified with precision. At the point of LOE, the fish were removed from the fish arena and returned to control temperature and salinity where they rapidly recovered. All fish were fasted for 24 hours before the CT_max_ test and there was 100% survival after the tests.

### Statistical analysis

For statistical analysis, each individual counted as one observation (N=263), belonging to one of the four salinity treatments (coded 0, 0.5, 1, 3 ppt), one of the time-points (coded 0 for 2 hours and 1 for 10 days), and one replicate (coded 1 or 2). To test for an effect of salinity and time-in-treatment on CTmax, we fitted a linear mixed effect model using CTmax (°C) as a response variable, salinity treatment and time-in-treatment as explanatory variables, and the salinity-time-replicate combination as a random effect on the intercept. F and p values were calculated using Satterthwaite’s method with type II tests. The mixed model was fitted using restricted maximum likelihood. All analysis was performed using *R* v4.2.1 (R Core Team, 2021) via *Rstudio* v2022.7.0.548 (RStudio Team, 2022). R packages used were *lme4* v1.1.30 (Bates et al., 2021) for mixed model analysis and *ggplot* v3.3.6 (Wickham, 2016) for plotting; as well as the *tidyverse* v1.3.1 (Wickham et al., 2019) package collection for general data management.

## RESULTS AND DISCUSSION

There was no detectable difference in CT_max_ between salinity treatments at either time point, and there was no difference between time points (Fig 2). A linear mixed effect model could detect no effect of salinity (β=0.03, SE=0.02) time (β=0.07, SE=0.08), or their interaction (β=-0.03, SE=0.03)(Suppl. table 2).

**Figure 1.**
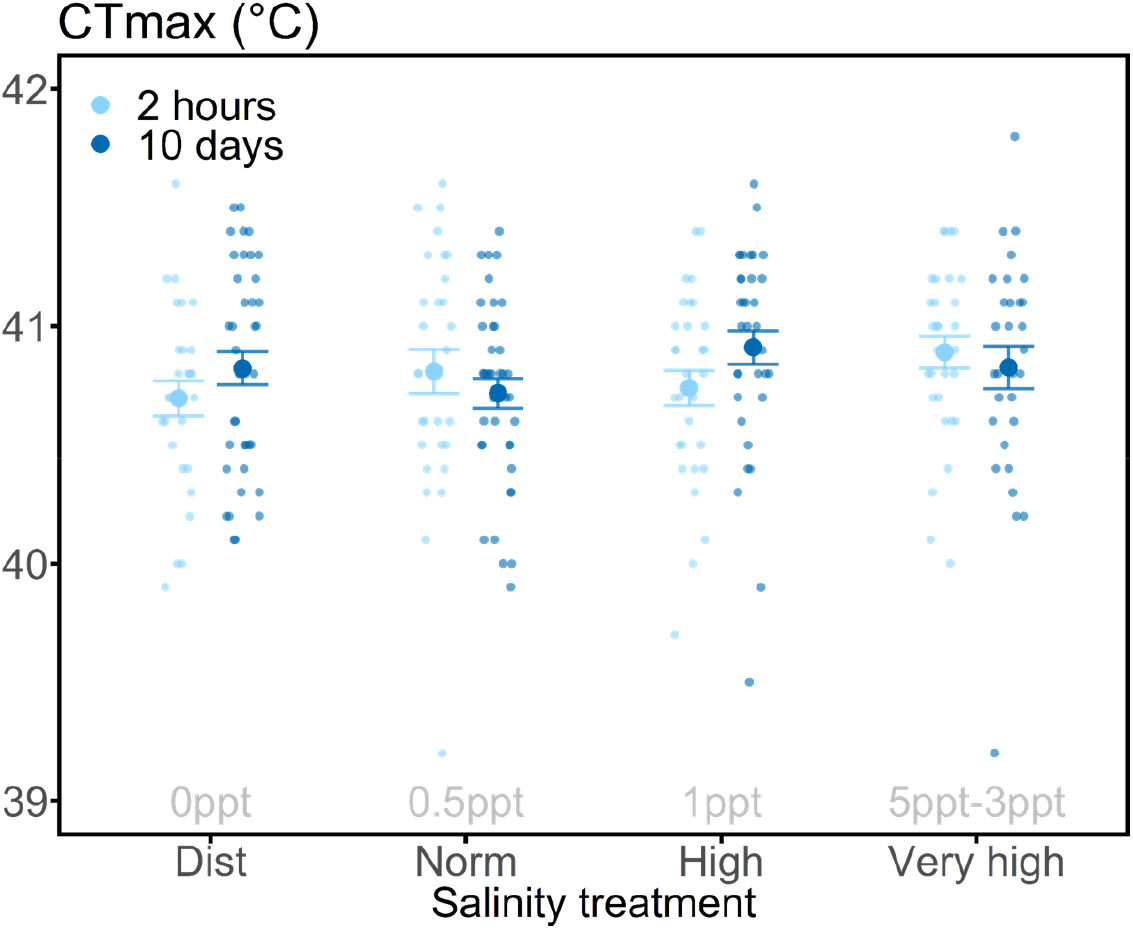
CT_max_ for individuals (N=263) tested at four different salinity treatments and two treatment durations (after 2 hours and after 10 days). Mean group results and SE is indicated by filled circles and error bars. Non-filled circles indicate CT_max_ of single individuals. Points are horizontally jittered for readability.

In the highest salinity treatment (3-5 ppt), five (of 69) individuals died during the experiment; These deaths happened at days three (n=1), seven (n=1) and ten (n=3). Furthermore, six individuals of the highest salinity treatment were removed from the CT_max_ test before reaching CT_max_ because they displayed abnormal behaviour during heating (described below). Two individuals from the 0 ppt 2.5hr treatment jumped out of the CT_max_ test tank and were excluded from the analysis. Mean CT_max_ for all individuals was 40.8 ± 0.03°C (Mean ± SE, N=263). The highest salinities used are 6 and 10 times higher than normal holding salinities for zebrafish (0.5 ppt), and 70 times higher than what has been observed in the wild for this species (0.06 ppt), and although the treatment appeared to affect the survival of the fish there was no detectable effect on acute thermal tolerance.

During CT_max_ testing of fish from the highest salinity treatment (at two weeks), an unexpected behaviour occurred in which fish would sometimes perform a sudden burst in swimming, followed by a cramped seizure-like spasm lasting a few seconds that caused loss of equilibrium. This happened at temperatures down to 10°C lower than the normal CT_max_ and was first mistaken as their final LOE at CT_max,_ hence the premature removal of 6 fish before their true CT_max_. This happened to several more individuals, and we found that fish always recovered after these cramping events and would continue swimming until reaching the normal LOE reaction at similar temperatures to other individuals. The total number of individuals this happened to was not recorded, but it only occurred in the highest salinity treatment after two weeks of exposure. This suggests that the highest salinity treatment had an effect on the tested fish, but didn’t affect thermal tolerance.

Our results paint a complicated picture of the interactions between salinity and thermal tolerance. On one side, the salinity increase of 3 ppt appeared high enough to elicit negative effects in Zebrafish; The physiological background of this effect is unclear, but the effect was enough to increase mortality and to trigger a temporary seizure during rapid warming in some fish. Based on this, we expect a salinity as high as 3 ppt to have a negative fitness effect on zebrafish, and these observations indicate that a high temperature does somehow interact with water salinity in a negative sense. On the other side, water salinity does not seem to interact with CT_max_, indicating that water salinity does not significantly interact with the physiological mechanisms determining acute upper thermal tolerance. We interpret the lack of effect of water salinity on acute thermal tolerance as a sign that these stressors have disparate effects acting on separate physiological and biochemical mechanisms. While the presence of an effect could have suggested a mechanistic connection between the salinity and heat stress, the observed lack of effect can be interpreted as evidence against such a connection.

Zebrafish were previously reported to show acute thermal tolerance that correlated negatively with habitat water salinity in a survey of wild zebrafish populations (Morgan et al., 2019; Sundin et al., 2019). That result does not match the observed robustness of CT_max_ to even larger salinity differences in the current study. We believe the present result is more reliable due to the higher sample size and broad range of salinities. The discrepancy may be due to an additional factor(s), associated with stream salinity and CT_max,_ confounding the correlation observed in Morgan et al. (2019), but we have no suggestions on what the confounding factor(s) might be.

To get a better overview of the relationship between salinity and CT_max_ in previous studies on this topic, we compiled a list of published papers where water salinity was manipulated experimentally, and where CT_max_ was tested as the outcome variable (Figure 2. Suppl. table 1). Compared to these previous studies, our study adds to the list of studies showing no effect, together with Davis et al. (2019) and Hines et al. (2019). A notable difference between the studies showing an effect and those not showing one is the salinity tolerance of the species used. This study and the study by Hines et al. (2019) used relatively low water salinities (up to 5 ppt and 12 ppt), while studies reporting an effect (Haney & Walsh, 2003; King & Sardella, 2017; Metzger et al., 2016; Sardella et al., 2008; Shaughnessy & McCormick, 2018) used far higher ranges (up to 25 ppt, and up to 60 ppt in one study). The studies showing an effect all used highly euryhaline species and acclimation times of at least one week, indicating that the effects on thermal tolerance may be connected to salinity acclimation itself or through some other general stress response. Still, the study by Hines et al. (2019) found no effect despite using salinities up to 30 ppt and acclimation times up to 600 days. Overall, looking at the published literature on the subject, it looks like the association between salinity and thermal tolerance is highly complex and species- and context-dependent.

## CONCLUSION

We show that water salinity does not affect the critical thermal maximum of zebrafish. However, zebrafish at high salinities (3 ppt) did show abnormal behaviour during the CT_max_ trial, indicating some interaction between warming- and salinity effects.

## ACKNOWLEDGEMENTS

We acknowledge and thank Miriam Dørum and Jan Arvid Sand for help with feeding and maintenance of the experimental setup.

## SUPPLEMENTARY MATERIAL

**Suppl. table 1.** Overview of studies reporting tests of CT_max_ under different salinity regimes. Shows for each study the CT_max_ ramping rate (R. rate), the range of acclimation temperatures used (Accl. temp. range), the range of salinities used (Sal. range), the salinity ranges where these species are observed in the wild (Nat. sal. Range), how long the fish got time to acclimate to their water salinities before testing CT_max_ (Sal. accl. time), and what effect increasing salinity had on CT_max_ (Effect on CT_max_). Some natural salinity ranges are gathered from independent studies (sources noted below table):

| Study/Species | CTmax r. rate | Accl. temp.range | Sal. range | Nat. sal. range | Sal. accl. time | Effect on CTmax |
| --- | --- | --- | --- | --- | --- | --- |
| **Shaughnessy & McCormick (2018)** |  |  |  |  |  |  |
| Brook trout (*Salvelinus fontinalis*) | 4 - 2°C/h | 18°C | FW - 25 ppt | 0-32 ppt<sup>(1)</sup> | 0, 2, 5, 16d | Reduced |
| Decreased CTmax during acclimation after acute transfer to seawater, but restored to normal after 16 days. CTmax ramping rate 4°C/h first hour, then 2°C/h. FW = freshwater (salinity not specified). |  |  |  |  |  |  |
| **Morgan et al (2019)** |  |  |  |  |  |  |
| Zebrafish (*Danio rerio*) | 0.3°C/min | 25-35°C | 0.03-0.10 ppt | 0.1-0.6 ppt<sup>(2)</sup> | Native | Reduced |
| Ctmax was negatively correlated with the water salinity of streams in wild zebrafish in India. |  |  |  |  |  |  |
| **Åsheim et al (this paper)** |  |  |  |  |  |  |
| Zebrafish (*Danio rerio*) | 0.3°C/min | 26°C | 0-5 ppt | 0.1-0.6 ppt<sup>(2)</sup> | 0, 10d | No effect |
| **Hines et al (2019)** |  |  |  |  |  |  |
| Atlantic salmon (*Salmo salar*) | 0.1°C/min | 12°C | 2.5-30 ppt | 0-14 ppt<sup>(3)</sup> | 60, 400d | No effect |
| Coho salmon (*Oncorhynchus kisutch*) | 0.1°C/min | 12°C | 2.5-30 ppt | 0-28 ppt<sup>(4)</sup> | 60, 400d | No effect |
| Tested 60 and 400 days post-smoltification. |  |  |  |  |  |  |
| **Davis et al (2019)** |  |  |  |  |  |  |
| Delta smelt (*Hypomesus transpacificus*) | 0.3°C/min | 16-20°C | 2.4-12 ppt | 0-18 ppt | 0, 2, 4, 7d | No effect |
| Mississippi Silverside (*Menidia Berylinna*) | 0.3°C/min | 16-20°C | 2.4-12 ppt | 0-35 ppt | 0, 2, 4, 7d | No effect |
| Largemouth bass (*Micropterus salmoides*) | 0.3°C/min | 16-20°C | 2.4-8 ppt | 0-4 ppt | 0, 2, 4, 7d | No effect |
| **Haney & Walsh (2003)** |  |  |  |  |  |  |
| Blackbanded limia (*Limia melanotata*) | 0.3°C/min | 25-35°C | 0-60 ppt | 0-70 ppt | 7-14d | Peak 30 ppt |
| **Sardella, Sanmarti & Krütz (2008)** |  |  |  |  |  |  |
| Green sturgeon (*Acipenser medirostris*) | 0.3°C/min | 18°C | 1-24 ppt | 8.8-32.1 ppt<sup>(5)</sup> | 7d | Peak 10 ppt |
| **King & Sardella (2017)** |  |  |  |  |  |  |
| Mozambique tilapia (*Oreochromis mossambicus*) | 0.3°C/min | 20-37 °C | 3-24 ppt | 0-120 ppt<sup>(6)</sup> | 28d | Increased* |
| *24g/l acclimated fish had a larger thermal tolerance window. |  |  |  |  |  |  |
| **Metzger, Healy, Schulte (2016)** |  |  |  |  |  |  |
| Three-spined Stickleback (*Gasterosteus acceatus*) | 0.3°C/min | 18°C | 2-20 ppt | 0-30 ppt<sup>(7)</sup> | Native, 14d | Increased |
| Two populations (freshwater, seawater), tested at both salinities. Fish acclimated to higher salinity had higher Ctmax. Salinity acclimation affected expression of heat shock proteins during heat shock test. |  |  |  |  |  |  |

References for natural salinity ranges are 1: (McCormick et al., 1998), 2: (Spence et al., 2006), 3: (Otto & McInerney, 1970), 4: (Halfyard et al., 2012), 5: (Israel & Klimley, 2008), 6: (Whitfield & Blaber, 1979), 7: (DeFaveri et al., 2013)

**Suppl. table table 2.**
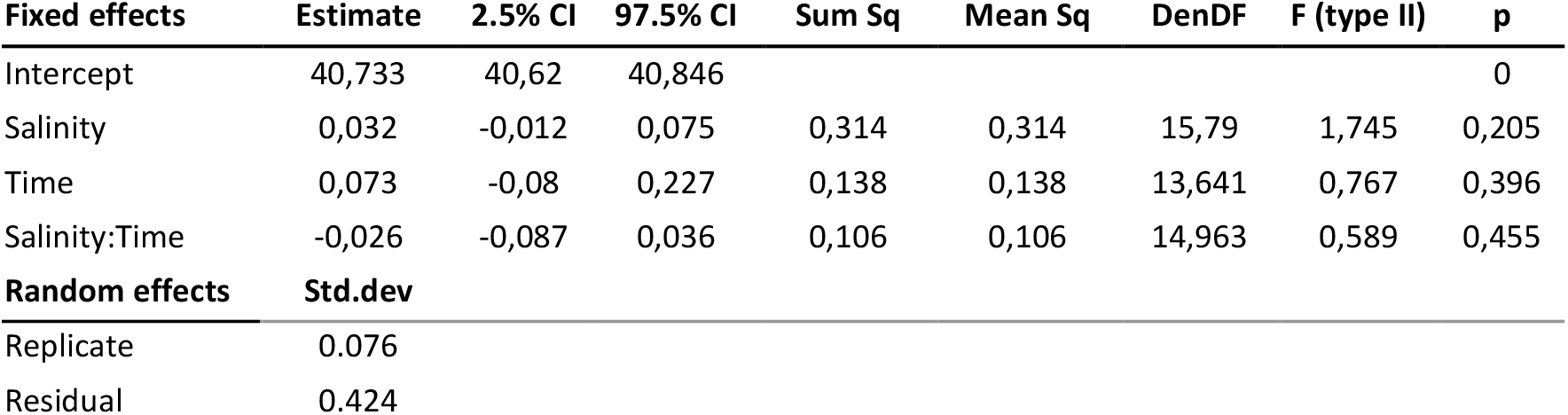
Statistical summary for the mixed-effect model on CT_max._ F and p values were calculated using Satterthwaite’s method with type II tests. The mixed model was fit using restricted maximum likelihood. CT_max_ was modelled to be linearly affected by salinity (1-5 ppt), time of exposure (2 hours or 10 days, coded 0 or 1), as well as an interaction between salinity and time. The estimated “Time” effect indicates the effect of a 10d exposure to the salinity treatment before testing CT_max_ (as opposed to 2 hours), and the “Salinity:Time” interaction indicates the additional change in ctmax for each increase in salinity at the 10d exposure time. No significant effect was found for any of the included factors.

## References

Bates, D., Maechler, M., Bolker, B., Walker, S., Christensen, R. H. B., Singmann, H., Dai, B., Scheipl, F., Grothendieck, G., Green, P., Fox, J., Bauer, A., & Pavel N. K. (2021). lme4: Linear Mixed-Effects Models using “Eigen” and S4 (1.1-27.1). https://CRAN.R-project.org/package=lme4

Becker, C. D., & Genoway, R. G. (1979). Evaluation of the critical thermal maximum for determining thermal tolerance of freshwater fish. Environmental Biology of Fishes, 4(3), 245. https://doi.org/10.1007/BF00005481

Davis, B. E., Cocherell, D. E., Sommer, T., Baxter, R. D., Hung, T.-C., Todgham, A. E., & Fangue, N. A. (2019). Sensitivities of an endemic, endangered California smelt and two non-native fishes to serial increases in temperature and salinity: Implications for shifting community structure with climate change. Conservation Physiology, 7(1), coy076. https://doi.org/10.1093/conphys/coy076

DeFaveri, J., Jonsson, P. R., & Merilä, J. (2013). Heterogeneous genomic differentiation in marine threespine sticklebacks: Adaptation along an environmental gradient: Special section. Evolution, 67(9), 2530–2546. https://doi.org/10.1111/evo.12097

Genin, A., Levy, L., Sharon, G., Raitsos, D. E., & Diamant, A. (2020). Rapid onsets of warming events trigger mass mortality of coral reef fish. Proceedings of the National Academy of Sciences, 117(41), 25378–25385. https://doi.org/10.1073/pnas.2009748117

Halfyard, E. A., Gibson, A. J. F., Ruzzante, D. E., Stokesbury, M. J. W., & Whoriskey, F. G. (2012). Estuarine survival and migratory behaviour of Atlantic salmon Salmo salar smolts. Journal of Fish Biology, 81(5), 1626–1645. https://doi.org/10.1111/j.1095-8649.2012.03419.x

Haney, D. C., & Walsh, S. J. (2003). Influence of salinity and temperature on the physiology of Limia melanonotata (Cyprinodontiforme: Poeciliidae): A search for abiotic factors limiting insular distribution in Hispaniola. Caribbean Journal of Science, 39(3), 327–337.

Hines, C. W., Fang, Y., Chan, V. K. S., Stiller, K. T., Brauner, C. J., & Richards, J. G. (2019). The effect of salinity and photoperiod on thermal tolerance of Atlantic and coho salmon reared from smolt to adult in recirculating aquaculture systems. Comparative Biochemistry and Physiology Part A: Molecular & Integrative Physiology, 230, 1–6. https://doi.org/10.1016/j.cbpa.2018.12.008

Israel, J. A., & Klimley, A. P. (2008). Life history conceptual model for North American green sturgeon (Acipenser medirostris). Delta Reg Ecosyst Restor Implement Plan Rep, 1–49.

King, M., & Sardella, B. (2017). The effects of acclimation temperature, salinity, and behavior on the thermal tolerance of Mozambique tilapia (Oreochromis mossambicus). Journal of Experimental Zoology Part A: Ecological and Integrative Physiology,327(7), 417–422. https://doi.org/10.1002/jez.2113

Lawrence, C. (2007). The husbandry of zebrafish (Danio rerio): A review. Aquaculture, 269(1), 1–20. https://doi.org/10.1016/j.aquaculture.2007.04.077

Lutterschmidt, W. I., & Hutchison, V. H. (1997). The critical thermal maximum: History and critique. Canadian Journal of Zoology, 75(10), 1561–1574. https://doi.org/10.1139/z97-783

McCormick, S. D., Hansen, L. P., Quinn, T. P., & Saunders, R. L. (1998). Movement, migration, and smolting of Atlantic salmon (Salmo salar). Canadian Journal of Fisheries and Aquatic Sciences, 55(S1), 77–92. https://doi.org/10.1139/d98-011

Metzger, D. C. H., Healy, T. M., & Schulte, P. M. (2016). Conserved effects of salinity acclimation on thermal tolerance and hsp70 expression in divergent populations of threespine stickleback (Gasterosteus aculeatus). Journal of Comparative Physiology B, 186(7), 879–889. https://doi.org/10.1007/s00360-016-0998-9

Morgan, R., Andreassen, A. H., Åsheim, E. R., Finnøen, M. H., Dresler, G., Brembu, T., Loh, A., Miest, J. J., & Jutfelt, F. (2022). Reduced physiological plasticity in a fish adapted to stable temperatures. Proceedings of the National Academy of Sciences, 119(22), e2201919119. https://doi.org/10.1073/pnas.2201919119

Morgan, R., Finnøen, M. H., & Jutfelt, F. (2018). CTmax is repeatable and doesn’t reduce growth in zebrafish. Scientific Reports, 8(1), 7099. https://doi.org/10.1038/s41598-018-25593-4

Morgan, R., Sundin, J., Finnøen, M. H., Dresler, G., Vendrell, M. M., Dey, A., Sarkar, K., & Jutfelt, F. (2019). Are model organisms representative for climate change research? Testing thermal tolerance in wild and laboratory zebrafish populations. 7. https://doi.org/10.1093/conphys/coz036

Otto, R. G., & McInerney, J. E. (1970). Development of Salinity Preference in Pre-Smolt Coho Salmon, Oncorhynchus kisutch. Journal of the Fisheries Research Board of Canada, 27(4), 793–800. https://doi.org/10.1139/f70-081

R Core Team. (2021). R: A Language and Environment for Statistical Computing. R Foundation for Statistical Computing. https://www.R-project.org/

RStudio Team. (2022). RStudio: Integrated Development Environment for R. RStudio, PBC. http://www.rstudio.com/

Sardella, B. A., Sanmarti, E., & Kültz, D. (2008). The acute temperature tolerance of green sturgeon (Acipenser medirostris) and the effect of environmental salinity. Journal of Experimental Zoology Part A: Ecological Genetics and Physiology, 309A(8), 477–483. https://doi.org/10.1002/jez.477

Seneviratne, S. I., Donat, M. G., Mueller, B., & Alexander, L. V. (2014). No pause in the increase of hot temperature extremes. Nature Climate Change, 4(3), 161–163. https://doi.org/10.1038/nclimate2145

Shaughnessy, C. A., & McCormick, S. D. (2018). Reduced thermal tolerance during salinity acclimation in brook trout (Salvelinus fontinalis) can be rescued by prior treatment with cortisol. Journal of Experimental Biology, jeb.169557. https://doi.org/10.1242/jeb.169557

Spence, R., Fatema, M. K., Reichard, M., Huq, K. A., Wahab, M. A., Ahmed, Z. F., & Smith, C. (2006). The distribution and habitat preferences of the zebrafish in Bangladesh. Journal of Fish Biology, 69(5), 1435–1448. https://doi.org/10.1111/j.1095-8649.2006.01206.x

Sundin, J., Morgan, R., Finnøen, M. H., Dey, A., Sarkar, K., & Jutfelt, F. (2019). On the Observation of Wild Zebrafish (Danio rerio) in India. Zebrafish, 16(6), 546–553. https://doi.org/10.1089/zeb.2019.1778

Wegner, K. M., Kalbe, M., Milinski, M., & Reusch, T. B. (2008). Mortality selection during the 2003 European heat wave in three-spined sticklebacks: Effects of parasites and MHC genotype. BMC Evolutionary Biology, 8(1), 124. https://doi.org/10.1186/1471-2148-8-124

Whitfield, A. K., & Blaber, S. J. M. (1979). The distribution of the freshwater cichlid Sarotherodon mossambicus in estuarine systems. Environmental Biology of Fishes, 4(1), 77–81. https://doi.org/10.1007/BF00005931

Wickham, H. (2016). ggplot2: Elegant Graphics for Data Analysis. Springer-Verlag New York. https://ggplot2.tidyverse.org

Wickham, H., Averick, M., Bryan, J., Chang, W., McGowan, L. D., François, R., Grolemund, G., Hayes, A., Henry, L., Hester, J., Kuhn, M., Pedersen, T. L., Miller, E., Bache, S. M., Müller, K., Ooms, J., Robinson, D., Seidel, D. P., Spinu, V., … Yutani, H. (2019). Welcome to the tidyverse. Journal of Open Source Software, 4(43), 1686. https://doi.org/10.21105/joss.01686

